# Fine-scale spatial patterns in hot springs mat bacterial communities

**DOI:** 10.1101/2025.03.11.642739

**Authors:** José Ignacio Arroyo, Carlos Pedrós-Alió, Martin F. Polz, Roy Mackenzie, Pablo A. Marquet, Beatriz Díez

## Abstract

Some of the most widely recognized spatial scaling relationships in ecology include the species-area, the abundance-occupancy, and the community compositional similarity-geographic distance relationships. These patterns have been shown to emerge from common mechanisms, such as habitat heterogeneity and colonization-extinction dynamics, and are predicted by various ecological models and theories, including Island Biogeography, Metapopulation theory, and Neutral Theory of Biodiversity and Biogeography. These patterns have been studied in microbial communities at large-scale, but studies at fine grain and small extent are rare, despite at a fine-scale, emerging patterns can be tested with a higher resolution. In this study, we explored these macroecological patterns in bacterial communities inhabiting hot springs mats, which have been shown to exhibit fine-scale spatial heterogeneity in fundamental environmental parameters such as temperature. In three different localities, we sampled a grid with a heterogeneous temperature at a fine-scale, a small extent (approximately 150 cm^2^), and fine grain (each containing 30 cells spaced about 1 or 2 cm apart). Our findings revealed sublinear scaling for the species-area relationship, with similar parameters across localities, indicating a low rate of spatial species turnover. For the abundance-occupancy relationship, we observed increasing trends, meaning that species that occupied more patches were, on average, more abundant. Additionally, we identified a decay in community compositional similarity with distance in two of the localities, though with low parameter values, indicating minimal geographic isolation at this scale. These results are consistent with different models that predict that spatial ecological patterns arise from spatial heterogeneity, as different species can partition their niches in space and constitute an example of predictable spatial patterns at a fine scale.

## Introduction

Understanding the ecological mechanisms driving the emergence of predictable spatial patterns in communities is a fundamental question in ecology (Levin, 1992). Several properties of ecological systems change with spatial scale, including population occupancy or occurrence (Harte et al., 2005; Kunin, 1998), abundance (Hanski & Gyllenberg, 1997), species diversity (Hanski & Gyllenberg, 1997), phylogenetic diversity (Morlon et al., 2011), functional diversity (Alirezazadeh et al., 2021), spatial covariance or correlation of populations (Bjørnstad et al., 1999; Liebhold et al., 2004), and the number of species interactions in an ecological network (Brose et al., 2004), among others. Various models and theories have been proposed to explain the origins of these patterns, some of which predict more than one of them. For instance, three interrelated theories have been developed to explain spatial patterns: Island Biogeography (MacArthur & Wilson, 2001), Metapopulation theory (Hanski, 1998; Levins, 1969), and the Unified Neutral Theory of Biodiversity and Biogeography (UNTBB; Hubbell, 2001). Each of these theories has developed a mathematical framework to describe the processes of colonization and extinction as populations move between suitable habitat patches (Figure 1). Insular biogeography models species richness on islands and as the sampling area increases (Rosenzweig, 1995). The number of species on an island or location is determined by colonization and extinction. Colonization rates depend on the distance to a source population (e.g., a nearby mainland), while extinction rates decrease with larger island sizes. One key prediction of this model is the species-area relationship. Metapopulation theory, originally developed by Levins (Levins, 1969), models the dynamics of the proportion of occupied habitat patches through colonization and extinction processes (Hanski & Gyllenberg, 1997; Levins, 1969). Levin’s original model has been extensively expanded, with one notable extension for multiple species predicting both species-area and occupancy-abundance relationships within a unified framework (Hanski & Gyllenberg, 1997). The Hanski and Gyllengerg (1997) metacommunity model is a framework describing the dynamics of species occupying different patches of suitable habitats or islands. This model applies to contexts such as mainland island systems or networks of multiple islands (Figure 1). It provides valuable insights into the interplay between colonization and extinction processes, offering a theoretical basis for understanding spatial scaling patterns in ecological systems. The UNTBB (Hubbell, 2001) models a metacommunity through stochastic birth-death processes, dispersal, and speciation and predicts the species-area and geographic distance-decay relationships under the assumption of neutrality.

**Figure 1.**
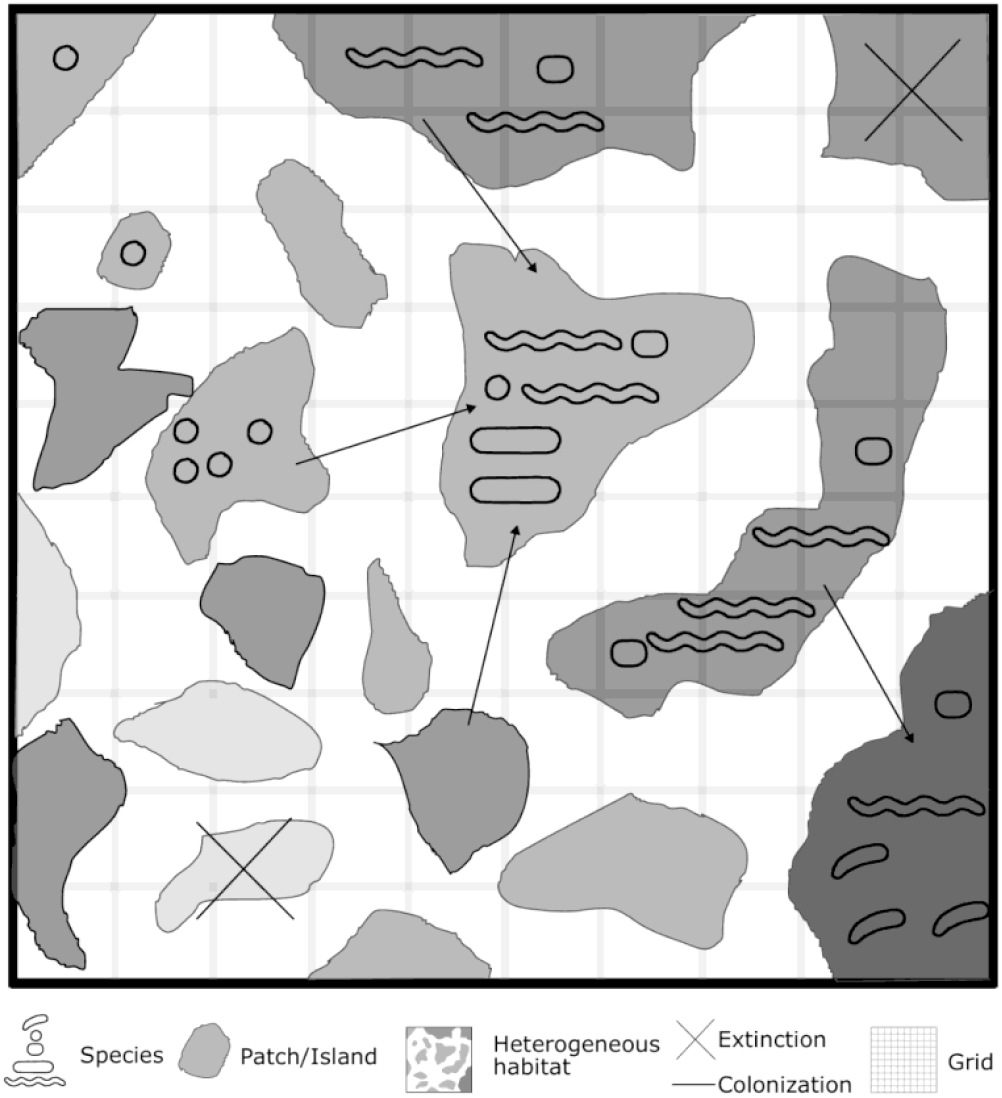
A metacommunity distributed in a heterogeneous landscape. A metacommunity consists of a set of populations of different species (represented by typical bacterial shapes), distributed in heterogeneous suitable habitat patches (represented by gray areas). The spatial structure of populations in such a heterogeneous landscape change through processes of colonization and the extinction of patches, as modeled in (Hanski & Gyllenberg, 1997), for example. Some of the emergent patterns of these processes are a positive relationship between the sampled area and the number of species and a positive relationship between the average abundance of a given species and the number or proportion of patches that the species occupies. Often, these two patterns are studied by dividing the landscape as a lattice or grid consisting of a set of quadrants of a given size (as depicted in the figure) and taking an increasing number of cells or areas from a given corner.

These three theories outlined above make several predictions, including the following three key patterns (Figure 2):

**Figure 2.**
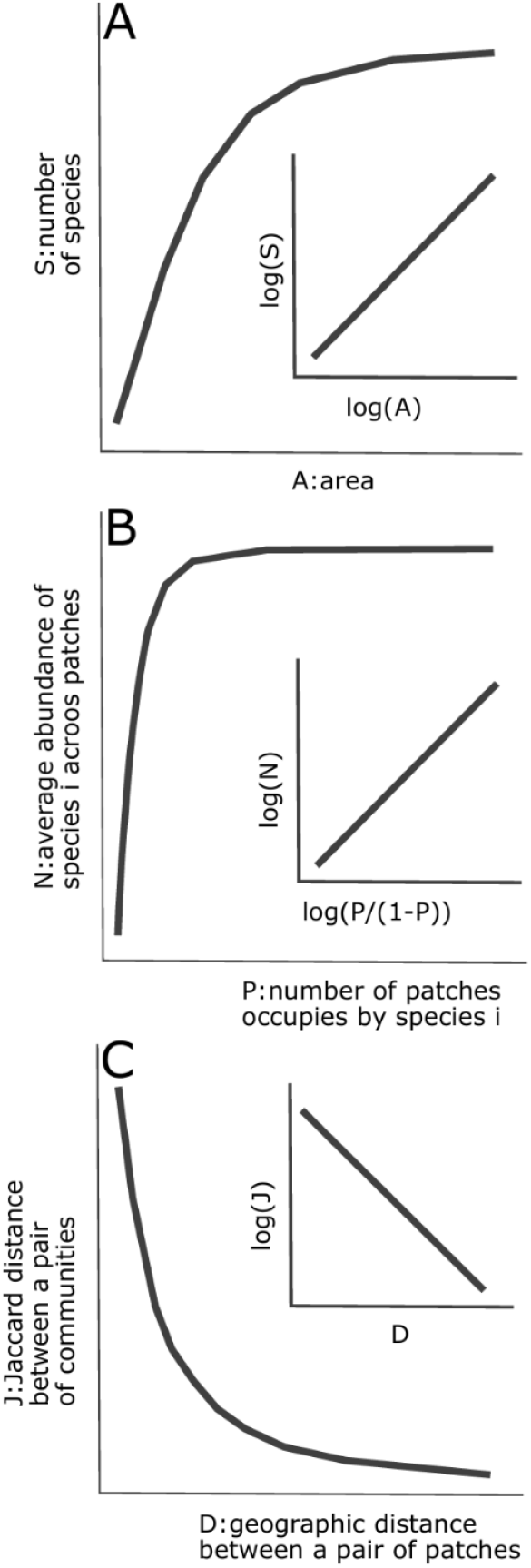
Model predictions for spatial patterns. Predicted patterns from models of dynamics of colonization and extinction through heterogeneous environments. A) species-area, B) abundance-occupancy, and C) compositional similarity-distance. The insets represent the linearized forms of the relationships. See equations 1-7.

i. Abundance-occupancy (or occurrence) relationship (e.g., the number of cells in a lattice) (Gaston et al., 2000; Hanski & Gyllenberg, 1997) (Figure 2A): Refers to the relationship between the abundance of individuals in an assemblage (a group of populations) across a set of localities and the number of those localities where the assemblage is present. Typically, it is studied by defining a lattice with a cell of arbitrary size (e.g., 1 m^2^) and counting the number of individuals within increasing sets of cells containing the assemblage. For example, one cell is selected, and the number of individuals is counted; then, two contiguous cells are combined, and the total abundance is summed, and so on. The pattern is characterized by an asymptotic increase in total abundance with the number of occupied cells. It is important to note that often, this pattern is expressed as occupancy-abundance, which is a more natural expression when deriving a model for these quantities, but abundance-occupancy is more useful for predictions.
ii. Species-area relationship (Hanski & Gyllenberg, 1997; Pan, 2013; Scheiner, 2003) (Figure 2B): This pattern refers to the increase in species richness with the size of the sampled area. It is often constructed by defining a lattice or plot, aggregating contiguous cells, and counting the accumulative number of species. Alternatively, it can be calculated as an average across sampled cells.
iii. Community similarity and geographic distance (distance-decay) (Figure 2C): This pattern describes the decline in compositional similarity (e.g., Jaccard index) between pairs of local communities as geographic distance increases. Predicted by several metacommunity models (Azaele et al., 2009; Chave & Leigh, 2002, 2002; Morlon et al., 2008), this pattern arises from colonization and extinction dynamics in heterogeneous patches.

Spatial scaling theories and models were originally conceived for macroorganisms, whose dispersal is often constrained by geographic barriers such as mountains or water bodies. Over the last decade, it has been investigated whether these theories also apply to microorganisms, despite their ecological processes operating at much smaller spatial scales. Most bacteria range in size from 1 to 10 micrometers, and their ecological interactions occur on a micrometer scale (Cordero & Datta, 2016; Heim et al., 2017; Ladau & Eloe-Fadrosh, 2019). Several field studies have examined spatial patterns at fine scales (Carreira et al., 2015; Lear et al., 2014; Youssef et al., 2010). For example, studies have explored environmental variation and its effects on microbial processes at scales ranging from centimeters to millimeters (O’Brien et al., 2016; Raynaud & Nunan, 2014; Vos et al., 2013). These works have shown the existence of patterns in community structure within a few centimeters, driven by factors such as topography, soil variables, and heavy metal contamination (Vos et al., 2013). At sub-millimeter scales (micrometers), environmental heterogeneity continues to influence microbial processes, but population interactions become increasingly significant (Bailey et al., 2013). However, while fine-scale variation in microbial communities has been reported, studies assessing whether these patterns agree with ecological theories are scarce (see Xue et al., 2021). The concept of scale is associated with the concept of extent (the size of the sampled area, which could be small or large) and grain (the size of the subunits of the sampled area, which can be fine or coarse) (Ladau & Eloe-Fadrosh, 2019; Wiens, 1989). Studies at a small scale are informative given that to a small extent, there is presumably little effect of dispersal limitation, and if samples are carried out at a fine grain the emerging pattern can be studied at high resolution.

Microbial mats in terrestrial hot springs provide a natural laboratory for studying small-scale ecological processes, given their small-scale heterogeneity. These ecosystems, are geothermally heated patches with temperatures higher than the mean environmental temperature (Pentecost et al., 2003). Compared to other terrestrial (e.g., soil) and aquatic ecosystems, hot spring microbial mats are less diverse (Lozupone & Knight, 2007) but contain representatives of most Archaeal and Bacterial lineages (Sharp et al., 2014). These communities can be found both as components of the pelagic environment in the water column or forming mats attached to the bottom sediments and other substrates. The latter are often more diverse than their pelagic counterparts (Beam et al., 2016).

Most studies examining ecological patterns in hot springs have been conducted at larger spatial scales (extents), often above the meter scale (Mackenzie et al., 2013; Podar et al., 2020; Sharp et al., 2014; Tamburello et al., 2022; Uribe-Lorío et al., 2019). At this large scale, community ecological patterns are explained by temperature (which can range from 10-160 ºC), pH, and other environmental factors. At a fine scale, despite spatial variation in environmental factors has been observed (e.g. Dunckel et al., 2009; Walther, K, 2013), but it has not been studied if the spatial ecological patterns agree with predictions made by ecological theories such as those described above. Testing ecological theories is relevant for at least two reasons, compared to a simple descriptive or comparative approach. Theories provide a fundamental understanding of mechanisms, and given their quantitative framework, they are useful to make predictions (Marquet et al., 2014).

In this study, for the first time, we assessed the existence of the macroecological patterns species-area, occupancy-abundance, and distance-decay relationships in hot springs microbial biofilms at fine spatial scales, considering both grain (sampling resolution) and extent (study area size, approximately 30 cm^2^ lattices) across three localities in Northern Patagonia. We identified consistent predictable patterns across all three localities. Furthermore, we observed distance-decay relationships and in two of them a relationship with temperature. Our study demonstrates that the processes of colonization and extinction operate at fine spatial scales in microbial communities, supporting the broader applicability of ecological scaling theories to microorganisms.

## Methods

### Sampling and data

We collected microbial mat samples from three hot springs in Northwestern Patagonia, specifically in the Comau fjord, Chile, during the summer (March 2011). The sampling sites included Cahuelmó (42º 15’ 11.8’’S / 72º 22’ 4.4’’W), Porcelana Geyser (referred to as Geyser from now on for simplicity; 42º 24’ 51’’S / 72º 29’ 2.2’’ W) and Porcelana hot spring (referred to as Porcelana; 42º 27’ 29.1’’ S – 72º 27’ 39.3’’ W) (Supplementary Figure S1). Cahuelmó is located 20.3 km northeast of Geyser and 24.0 km from Porcelana, on the eastern side of Comau Fjord (Supplementary Figure 1).

Geyser and Porcelana hot springs are situated on the western side of the fjord, within the Huequi Peninsula, approximately 5.2 km apart. In each of these three locations, we collected 30 fine-scale samples within a grid-like plot (Supplementary Figure 21). Simultaneously during sampling, pH was measured for each grid point using pH indicator strips, and the temperature was recorded using a digital thermometer (Oakton, model 35607-85). All sampled mat plots showed approximately neutral pH (near pH 6-7). For Cahuelmó and Geyser, we sampled grids measuring approximately 20 × 8 cm, each containing 30 circular samples of 1 cm diameter and 1 cm thick arranged in three rows per ten columns. At Porcelana, we sampled a microbial ecotone marking the boundary between two distinct microbial communities: an orange and a dark green one (Supplementary Figure 2). The grid at Porcelana measured approximately 15 × 10 cm in the horizontal plane of the microbial mat and included 30 circular samples, each 1 cm in diameter and approximately 2 cm thick, arranged in six rows per five columns. This sampling design ensured that all plots had approximately equal total areas (∼133 cm^2^), with each grid cell covering ∼5.33 cm^2^, and 25 cells were sampled per grid.

Each locality had a water source with different temperature ranges and surrounding environmental conditions (Mackenzie et al. 2013, Alcamán et al. 2015, Alcorta et al. 2018). Within the sampled grids, the temperature varied as follows: Cahuelmó 31.1 - 44.7 (a 13.6 °C range), Geyser 32.8 - 53.7 (a 20.9°C range), and Porcelana 52.4 - 60.2 (a 7.8°C range) (Supplementary Figure 2). These temperature ranges correspond to what we refer to as low (Cahuelmó), intermediate (Geyser), and high (Porcelana) temperature localities. Distinct spatial patterns of temperature variation were observed, with temperature differences increasing as the sampled area expanded. For simplicity, we will refer to the analyses of patterns within a single grid at a locality as small-scale or fine-scale. While this sampling design was not specifically intended to explore the effects of small temperature ranges, we briefly discuss analyses exploring the relationships between abundance, richness, and temperature.

Details regarding DNA extraction, library preparation, sequencing, pre-processing, analysis of reads, and statistical analysis are given in the Supplementary Materials. Taxonomic assignment and identification yielded a matrix of 549 operational taxonomic units (OTUs) across 80 samples. For subsequent analysis, this matrix was rarefied to 505 reads per sample, as implemented in the R (R Core Team) package vegan (Oksanen et al., 2017).

### Assessing the relationships

The relationships between the number of OTUs (alfa diversity) and the area were assessed according to

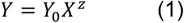

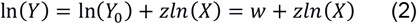

where Y0 is a constant and z is the scaling exponent (or slope in log scale). The relationship between the number of OTUs and area was determined by taking a spatially contiguous number of cells/patches and measuring the average number of OTUs in those patches. So first we took one cell and counted the number of OTUs, then we chose one contiguous patch to the already selected and averaged the number of OTUs in those two patches, then we chose the two patches and third contiguous one, i.e., three patches, and averaged the number of OTUs in those three patches, and so on until all patches are selected.

The interspecific abundance-occupancy, i.e., local average abundance and regional occupancy, was estimated as explained in (Borregaard & Rahbek, 2010; Gaston et al., 2000). The proportion of occupied cells was estimated as the number of occupied cells over the total number of cells in the grid, i.e. the maximum possible occupancy is 1. The abundance corresponds to the average abundance across all occupied cells and was calculated as average abundance=sum of abundances across all occupied cells/number of occupied cells.

We assessed the relationship between occupancy, estimated as proportional occupancy (P), and abundance following the suggestion made by (Hanski & Gyllenberg, 1997). These authors envisioned this relationship as logistic, which can be linearized by using a logit –transformation of the form:

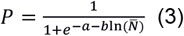

where p is the proportion of occupied patches in the grid and 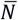is the average abundance of the OTU along all occupied patches. We can rearrange Eq. 3 to express abundance in terms of occupancy; 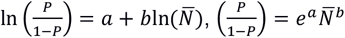,

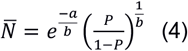

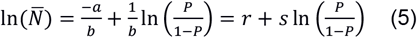

To quantify local community compositional similarity, we used the Jaccard index (J), as implemented in the function “betadiver” of the vegan package (Oksanen, F.J., et al., 2017) of the R environment (*R a Language and Environment for Statistical Computing*, 2010), and assessed its relationship against geographic distance (D), according to,

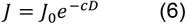

Where *J*_0_ and c are constants. For simplicity, we fitted the model in log scale,

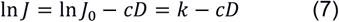

## Results

We analyzed three spatial patterns, the number of species and area, the average abundance of populations (OTUs) and the number of patches they occupy, and the compositional similarity and distance among pairs of patches, as predicted by equations 2, 5, and 7 (insets of Figure 2), in three fine-scale bacterial mats in hot springs. The results of these analyses are summarized in Table 1 and Figure 3. The species-area relationship was assessed by taking an increasing and spatially contiguous number of cells in the grid and averaging the number of OTUs in the included cells. When we scaled the average number of OTUs with the accumulated area, Cahuelmó and Porcelana showed similar slopes (varying between 0.31 to 0.43), while the Geyser locality displayed slightly different slopes (Figure 3).

**Table 1.** Estimated parameters, r-squared, and p-values for spatial scaling relationships fitted to each of the sites.

|  | Slope | Intercept | R<sup>2</sup> | P-value |
| --- | --- | --- | --- | --- |
| Number of OTUs-area (All) |  |  |  |  |
| Cahuelmó | 0.4 | 4.06 | 0.93 | <0.001 |
| Geyser | 0.31 | 3.95 | 0.97 | <0.001 |
| Porcelana | 0.43 | 4.18 | 0.97 | <0.001 |
| Abundance-occupancy (All) |  |  |  |  |
| Cahuelmó | 0.31 | 0.95 | 0.57 | <0.001 |
| Porcelana | 0.27 | 0.79 | 0.57 | <0.001 |
| Geyser | 0.27 | 0.92 | 0.55 | <0.001 |
| Jaccard-distance |  |  |  |  |
| Cahuelmó | -0.063 | -0.87 | 0.013 | 0.04 |
| Porcelana | -0.02 | -1.09 | 0.0016 | 0.48 |
| Geyser | -0.050 | -1.03 | 0.014 | 0.01 |

**Figure 3.**
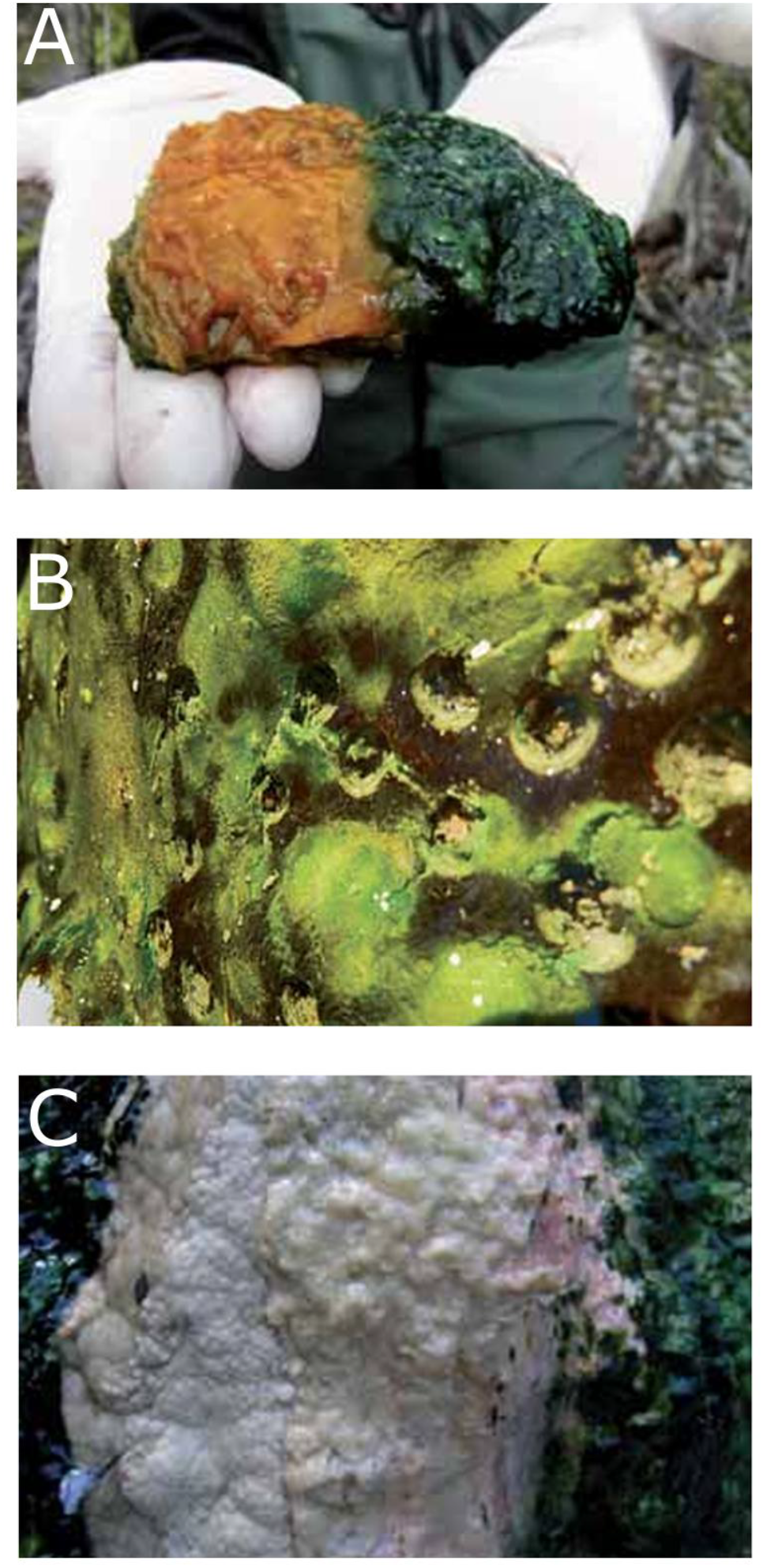
Photographs of the experimental design at small extent (e.g. panel A), and fine grain (e.g. panel B) samples. A) the Porcelana site exhibiting a small extent of approximately 10cmx15cm, B) the Cahuelmo site, exhibiting the 1cm holes separated by approximately 1 cm after sample extraction, and C) the Geyser site.

**Figure 4.**
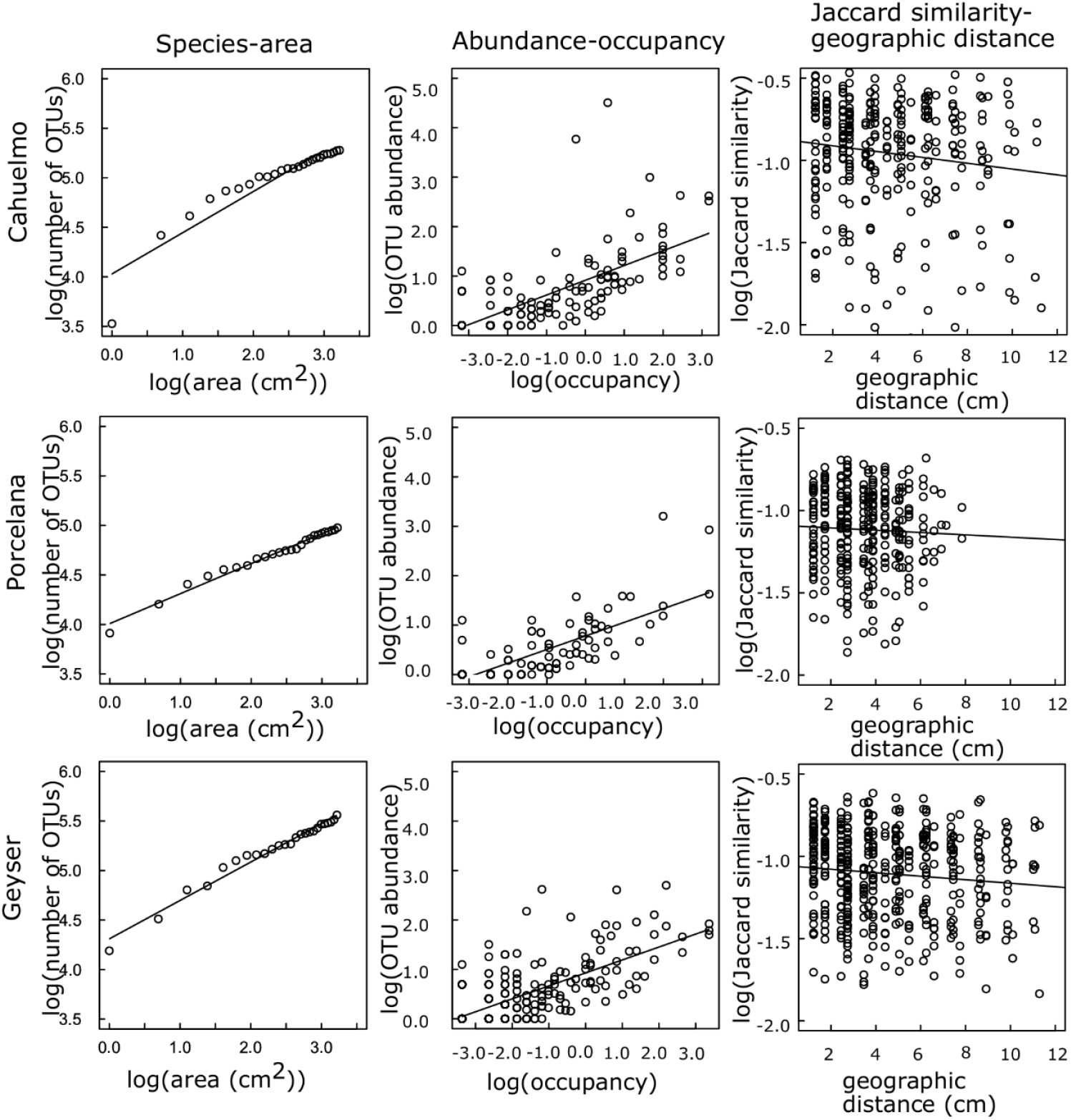
Spatial patterns at fine-scale bacterial mats. Each column is a different pattern, and each row is a different thermal spring. See Table 1 for estimated parameters and goodness-of-fit statistics.

We also explored the abundance-occupancy relationship for all OTUs. This relationship was constructed by counting the number of cells where a given OTU was present and averaging its abundance across these cells. Each point in this relationship corresponds to a unique OTU. In all cases, the abundance-occupancy relationship fitted well with Equation 5 and was similar (varying between 0.27 to 0.31) across the three patches (Figure 3). Additionally, the relationship between the compositional community similarity, measured using the Jaccard index, and geographic distance among pairs of patches was significant only in the Cahuelmó and Porcelana grids, in both cases exhibiting low exponents (varying between -0.02 to -0.06; Table 1, Figure 3), indicating a very low distance effect.

## Discussion

In this contribution, we showed that at spatial scales of just a few centimeters, it is possible to identify predictable ecological patterns. These include the increase in species richness with cumulative sampled area, the rise in average abundance of OTUs with occupied patches, and, at this scale, a minimal or zero decay in community composition similarity with distance. Such patterns can be explained through simple metacommunity models, where the proportion of occupied patches is influenced by colonization and extinction rates.

Understanding the predictability of fine-scale spatial patterns in microbial community structure is crucial for (at least) two key reasons. First, ecological theory has the potential to significantly advance microbial ecology (Marquet et al., 2014, Barberán, 2014; Prosser et al., 2007; Widder et al., 2016; Zhou, 2009) by validating or refining existing theories to explain patterns across both macro- and microorganisms. Second, understanding the characteristic scale of microorganisms-the scale at which patterns emerge (Levin, 1992)- is essential for designing effective sampling strategies in microbial ecology. Notably, the spatial parameters we identified at the centimeter scale for bacterial communities are comparable to those observed for macroorganisms but at larger scales (Durrett & Levin, 1996; Hanski & Gyllenberg, 1997). This suggests that the characteristic scale of microbial community dynamics operate at the centimeter scale.

Fine spatial or temporal scale microbial variation has also been documented in diverse environments, including soil (Carreira et al., 2015; O’Brien et al., 2016; Vos et al., 2013; Youssef et al., 2010) and aquatic ecosystems (Lear et al., 2014). These studies indicate that microbial communities operate at a scale much smaller than macroorganisms, where ‘local” scales encompass single sampling points and ‘regional” scales span only a few centimeters. However, the scale of variation is ecosystem-dependent, with aquatic systems often exhibiting greater connectivity (Sunagawa et al., 2015) than terrestrial ecosystems (Nemergut et al., 2013). It is also important to remark that ecosystems such as hots springs display fine-scale spatial heterogeneity at larger scales, where similar ranges of variation may be explained by the fractal structure of landscapes (Burrough, 1981).

A variety of models have been developed to explain all the different spatial scaling patterns observed in microbial communities and assessed in this study, among them the common approach involves the use of spatial Poisson point processes to model these patterns (Morlon et al., 2008; Picard & Favier, 2011; Plotkin et al., 2000). The species-area and abundance-occupancy relationships can often be predicted using models of random colonization and extinction, such as Island Biogeography or Metapopulation theories, and more recently, Neutral Theory. Some of these models are based in hypotheses that include habitat diversity, area per se, and passive sampling (Connor & McCoy 2001). Among these, environmental heterogeneity stands out as a key factor. Given the strong temperature heterogeneity in the bacterial mats studied, temperature likely plays a significant role in shaping these patterns. In mechanistic terms, it is hypothesized that habitats with higher environmental heterogeneity in fitness-related variables, such as temperature, may promote greater species abundance and diversity by enabling the coexistence of species with different niche breadths, ultimately meaning that areas with higher heterogeneity should have more species. This explains why increasing sampled area in heterogeneous environments results in higher species richness.

While our study is observational and not designed to differentiate among mechanisms, we hypothesize that both colonization-extinction dynamics and temperature heterogeneity contribute to the observed patterns. Additionally, we acknowledge that other chemical parameters, which were not measured in this study, could vary at centimeter scales and may also influence microbial spatial ecological patterns. The low geographic isolation observed at the centimeter scale in these environments suggests that the porous nature of microbial mats does not act as a significant geographic dispersal barrier to water flow. Consequently, populations can migrate across space, giving rise to emergent spatial scaling patterns. In other environments, centimeter-scale effects of distance have been documented (Yan et al., 2019), indicating that additional factors may influence the dispersion of microbial populations.

The spatial ecological patterns reported here raise several important questions. For instance, how can simple models, such as Hanski and Gyllenberg (1997), be extended to predict distance decay in microbial communities? Furthermore, how can independent modeling frameworks be integrated to generate new predictions? Several studies have explored these questions by employing a specific model or some from a single framework, with different mathematical forms (e.g., power laws, logarithmic models, etc.). Studies such as Hubbell’s Unified Neutral Theory of Biodiversity (2001) and Wang et al. (2009), which integrated the species-area relationship with the Metabolic Theory of Ecology to predict the temperature dependence of the exponent of the species-area relationships, illustrate the potential for integrating patterns into cohesive biodiversity theories. These are just a few examples of the potential to extend these frameworks into a more cohesive theory for biodiversity. In this sense, it is worth mentioning that there are other models and patterns than those studied here that describe the spatial scaping of a population (Harte et al., 2005; Hui et al., 2009; Hui & McGeoch, 2007) that we did not test here because we focused on communities. Among them are included the scaling of occupancy and area of a population, for example.

In conclusion, our findings, summed to evidence previously reported in other terrestrial and aquatic environments, highlight that microbial communities exhibit predictable patterns at fine spatial scales. These patterns are attributable to environmental spatial heterogeneity and, possibly, population interactions. From a practical perspective, our results highlight the importance of considering predictable spatial scaling patterns when designing microbial sampling strategies, as meaningful ecological trends may emerge within areas no larger than the size of a human hand.

## Acknowledgments

JIA was supported by a Beca de Doctorado Nacional granted by Agencia Nacional de Investigacion y Desarrollo (ANID) Grant 21130515. RM by Becas Chile (ANID) Grant 72100502. BD by the MILENIO ICN2021_044, FONDAP 1523A0002, and FONDECYT 1230217 from the ANID. CPA was supported by AYUDAS PARA LA REALIZACiÓN DE ESTANCIAS DE INVESTIGACiÓN EN EL CENTRO CIENTíFICO DE LA FUNDACiÓN HUINAY del CSIC 2011 y 2013. BD was supported by ANID – Millennium Science Initiative Program – ICN2021_044, Center for Climate and Resilience Research (CR)2, Chile FONDAP/ANID 1523A0002, and ANID-FONDECYT grants 1150171 and 1230217.

## Author contributions

RM and BD performed the sampling and carried out the OTU annotation. JIA performed the statistical and theoretical (data fitting) analysis. JIA, BD, CPA, MP, RM, and PM wrote the paper.

## References

Alirezazadeh, S., Borges, P. A. V., Cardoso, P., Gabriel, R., Rigal, F., & Borda-de-Água, L. (2021). Spatial Scaling Patterns of Functional Diversity. Frontiers in Ecology and Evolution, 9, 607177. 10.3389/fevo.2021.607177

Azaele, S., Muneepeerakul, R., Maritan, A., Rinaldo, A., & Rodriguez-Iturbe, I. (2009). Predicting spatial similarity of freshwater fish biodiversity. Proceedings of the National Academy of Sciences, 106(17), 7058–7062. 10.1073/pnas.0805845106

Bailey, V. L., McCue, L. A., Fansler, S. J., Boyanov, M. I., DeCarlo, F., Kemner, K. M., & Konopka, A. (2013). Micrometer-scale physical structure and microbial composition of soil macroaggregates. Soil Biology and Biochemistry, 65, 60–68. 10.1016/j.soilbio.2013.02.005

Barberán, A. (2014). The microbial contribution to macroecology. Frontiers in Microbiology, 5. 10.3389/fmicb.2014.00203

Beam, J. P., Bernstein, H. C., Jay, Z. J., Kozubal, M. A., Jennings, R. deM., Tringe, S. G., & Inskeep, W. P. (2016). Assembly and Succession of Iron Oxide Microbial Mat Communities in Acidic Geothermal Springs. Frontiers in Microbiology, 7. 10.3389/fmicb.2016.00025

Bjørnstad, O. N., Ims, R. A., & Lambin, X. (1999). Spatial population dynamics: Analyzing patterns and processes of population synchrony. Trends in Ecology & Evolution, 14(11), 427–432. 10.1016/S0169-5347(99)01677-8

Borregaard, M. K., & Rahbek, C. (2010). Causality of the Relationship between Geographic Distribution and Species Abundance. The Quarterly Review of Biology, 85(1), 3–25. 10.1086/650265

Brose, U., Ostling, A., Harrison, K., & Martinez, N. D. (2004). Unified spatial scaling of species and their trophic interactions. Nature, 428(6979), 167–171. 10.1038/nature02297

Burrough, P. A. (1981). Fractal dimensions of landscapes and other environmental data. Nature, 294(5838), 240–242. 10.1038/294240a0

Carreira, C., Piel, T., Staal, M., Stuut, J.-B. W., Middelboe, M., & Brussaard, C. P. D. (2015). Microscale spatial distributions of microbes and viruses in intertidal photosynthetic microbial mats. SpringerPlus, 4(1), 239. 10.1186/s40064-015-0977-8

Chave, J., & Leigh, E. G. (2002). A Spatially Explicit Neutral Model of β-Diversity in Tropical Forests. Theoretical Population Biology, 62(2), 153–168. 10.1006/tpbi.2002.1597

Connor, E. F., & McCoy, E. D. (2001). Species-area relationships. In Enciclopedia of biodiversity.

Cordero, O. X., & Datta, M. S. (2016). Microbial interactions and community assembly at microscales. Current Opinion in Microbiology, 31, 227–234. 10.1016/j.mib.2016.03.015

Dunckel, A. E., Cardenas, M. B., Sawyer, A. H., & Bennett, P. C. (2009). High-resolution in-situ thermal imaging of microbial mats at El Tatio Geyser, Chile shows coupling between community color and temperature. Geophysical Research Letters, 36(23), 2009GL041366. 10.1029/2009GL041366

Durrett, R., & Levin, S. (1996). Spatial Models for Species-Area Curves. Journal of Theoretical Biology, 179(2), 119–127. 10.1006/jtbi.1996.0053

Gaston, K. J., Blackburn, T. M., Greenwood, J. J. D., Gregory, R. D., Quinn, R. M., & Lawton, J. H. (2000). Abundance–occupancy relationships. Journal of Applied Ecology, 37(1), 39–59. 10.1046/j.1365-2664.2000.00485.x

Hanski, I. (1998). Metapopulation dynamics. Nature, 396(6706), 41–49. 10.1038/23876

Hanski, I., & Gyllenberg, M. (1997). Uniting Two General Patterns in the Distribution of Species. Science, 275(5298), 397–400. 10.1126/science.275.5298.397

Harte, J., Conlisk, E., Ostling, A., Green, J. L., & Smith, A. B. (2005). A THEORY OF SPATIAL STRUCTURE IN ECOLOGICAL COMMUNITIES AT MULTIPLE SPATIAL SCALES. Ecological Monographs, 75(2), 179–197. 10.1890/04-1388

Heim, N. A., Payne, J. L., Finnegan, S., Knope, M. L., Kowalewski, M., Lyons, S. K., McShea, D. W., Novack-Gottshall, P. M., Smith, F. A., & Wang, S. C. (2017). Hierarchical complexity and the size limits of life. Proceedings of the Royal Society B: Biological Sciences, 284(1857), 20171039. 10.1098/rspb.2017.1039

Hubbell, S. P. (2001). The unified neutral theory of biodiversity and biogeography (Nachdr.). Princeton Univ. Press.

Hui, C., & McGeoch, M. A. (2007). A self-similarity model for the occupancy frequency distribution. Theoretical Population Biology, 71(1), 61–70. 10.1016/j.tpb.2006.07.007

Hui, C., McGeoch, M. A., Reyers, B., Roux, P. C., Greve, M., & Chown, S. L. (2009). Extrapolating population size from the occupancy–abundance relationship and the scaling pattern of occupancy. Ecological Applications, 19(8), 2038–2048. 10.1890/08-2236.1

Kunin, W. E. (1998). Extrapolating Species Abundance Across Spatial Scales. Science, 281(5382), 1513–1515. 10.1126/science.281.5382.1513

Ladau, J., & Eloe-Fadrosh, E. A. (2019). Spatial, Temporal, and Phylogenetic Scales of Microbial Ecology. Trends in Microbiology, 27(8), 662–669. 10.1016/j.tim.2019.03.003

Lear, G., Bellamy, J., Case, B. S., Lee, J. E., & Buckley, H. L. (2014). Fine-scale spatial patterns in bacterial community composition and function within freshwater ponds. The ISME Journal, 8(8), 1715–1726. 10.1038/ismej.2014.21

Levin, S. A. (1992). The Problem of Pattern and Scale in Ecology: The Robert H. MacArthur Award Lecture. Ecology, 73(6), 1943–1967. 10.2307/1941447

Levins, R. (1969). Some Demographic and Genetic Consequences of Environmental Heterogeneity for Biological Control. Bulletin of the Entomological Society of America, 15(3), 237–240. 10.1093/besa/15.3.237

Liebhold, A., Koenig, W. D., & Bjørnstad, O. N. (2004). Spatial Synchrony in Population Dynamics. Annual Review of Ecology, Evolution, and Systematics, 35(1), 467–490. 10.1146/annurev.ecolsys.34.011802.132516

Lozupone, C. A., & Knight, R. (2007). Global patterns in bacterial diversity. Proceedings of the National Academy of Sciences, 104(27), 11436–11440. 10.1073/pnas.0611525104

MacArthur, R. H., & Wilson, E. O. (2001). The theory of island biogeography. Princeton University Press.

Mackenzie, R., Pedrós-Alió, C., & Díez, B. (2013). Bacterial composition of microbial mats in hot springs in Northern Patagonia: Variations with seasons and temperature. Extremophiles, 17(1), 123–136. 10.1007/s00792-012-0499-z

Marquet, P. A., Allen, A. P., Brown, J. H., Dunne, J. A., Enquist, B. J., Gillooly, J. F., Gowaty, P. A., Green, J. L., Harte, J., & Hubbell, S. P. (2014). On theory in ecology. BioScience, 64(8), 701– 710.

Morlon, H., Chuyong, G., Condit, R., Hubbell, S., Kenfack, D., Thomas, D., Valencia, R., & Green, J. L. (2008). A general framework for the distance–decay of similarity in ecological communities. Ecology Letters, 11(9), 904–917. 10.1111/j.1461-0248.2008.01202.x

Morlon, H., Schwilk, D. W., Bryant, J. A., Marquet, P. A., Rebelo, A. G., Tauss, C., Bohannan, B. J. M., & Green, J. L. (2011). Spatial patterns of phylogenetic diversity. Ecology Letters, 14(2), 141– 149. 10.1111/j.1461-0248.2010.01563.x

Nemergut, D. R., Schmidt, S. K., Fukami, T., O’Neill, S. P., Bilinski, T. M., Stanish, L. F., Knelman, J. E., Darcy, J. L., Lynch, R. C., Wickey, P., & Ferrenberg, S. (2013). Patterns and Processes of Microbial Community Assembly. Microbiology and Molecular Biology Reviews, 77(3), 342– 356. 10.1128/MMBR.00051-12

O’Brien, S. L., Gibbons, S. M., Owens, S. M., Hampton-Marcell, J., Johnston, E. R., Jastrow, J. D., Gilbert, J. A., Meyer, F., & Antonopoulos, D. A. (2016). Spatial scale drives patterns in soil bacterial diversity. Environmental Microbiology, 18(6), 2039–2051. 10.1111/1462-2920.13231

Oksanen, F.J., et al. (2017). Vegan: Community Ecology Package. R package Version 2.4-3. Https://CRAN.R-project.org/package=vegan.

Pan, X. (2013). Fundamental equations for species-area theory. Scientific Reports, 3(1), 1334. 10.1038/srep01334

Pentecost, A., Jones, B., & Renaut, R. W. (2003). What is a hot spring? Canadian Journal of Earth Sciences, 40(11), 1443–1446. 10.1139/e03-083

Picard, N., & Favier, C. (2011). A Point-Process Model for Variance-Occupancy-Abundance Relationships. The American Naturalist, 178(3), 383–396. 10.1086/661249

Plotkin, J. B., Potts, M. D., Leslie, N., Manokaran, N., Lafrankie, J., & Ashton, P. S. (2000). Species-area Curves, Spatial Aggregation, and Habitat Specialization in Tropical Forests. Journal of Theoretical Biology, 207(1), 81–99. 10.1006/jtbi.2000.2158

Podar, P. T., Yang, Z., Björnsdóttir, S. H., & Podar, M. (2020). Comparative Analysis of Microbial Diversity Across Temperature Gradients in Hot Springs From Yellowstone and Iceland. Frontiers in Microbiology, 11, 1625. 10.3389/fmicb.2020.01625

Prosser, J. I., Bohannan, B. J. M., Curtis, T. P., Ellis, R. J., Firestone, M. K., Freckleton, R. P., Green, J. L., Green, L. E., Killham, K., Lennon, J. J., Osborn, A. M., Solan, M., Van Der Gast, C. J., & Young, J. P. W. (2007). The role of ecological theory in microbial ecology. Nature Reviews Microbiology, 5(5), 384–392. 10.1038/nrmicro1643

R a language and environment for statistical computing: Reference index. (2010). R Foundation for Statistical Computing.

Raynaud, X., & Nunan, N. (2014). Spatial Ecology of Bacteria at the Microscale in Soil. PLoS ONE, 9(1), e87217. 10.1371/journal.pone.0087217

Rosenzweig, M. L. (1995). Species Diversity in Space and Time (1st ed.). Cambridge University Press. 10.1017/CBO9780511623387

Scheiner, S. M. (2003). Six types of species-area curves. Global Ecology and Biogeography, 12(6), 441–447. 10.1046/j.1466-822X.2003.00061.x

Sharp, C. E., Brady, A. L., Sharp, G. H., Grasby, S. E., Stott, M. B., & Dunfield, P. F. (2014). Humboldt’s spa: Microbial diversity is controlled by temperature in geothermal environments. The ISME Journal, 8(6), 1166–1174. 10.1038/ismej.2013.237

Sunagawa, S., Coelho, L. P., Chaffron, S., Kultima, J. R., Labadie, K., Salazar, G., Djahanschiri, B., Zeller, G., Mende, D. R., Alberti, A., Cornejo-Castillo, F. M., Costea, P. I., Cruaud, C., d’Ovidio, F., Engelen, S., Ferrera, I., Gasol, J. M., Guidi, L., Hildebrand, F., … Velayoudon, D. (2015). Structure and function of the global ocean microbiome. Science, 348(6237), 1261359. 10.1126/science.1261359

Tamburello, G., Chiodini, G., Ciotoli, G., Procesi, M., Rouwet, D., Sandri, L., Carbonara, N., & Masciantonio, C. (2022). Global thermal spring distribution and relationship to endogenous and exogenous factors. Nature Communications, 13(1), 6378. 10.1038/s41467-022-34115-w

Uribe-Lorío, L., Brenes-Guillén, L., Hernández-Ascencio, W., Mora-Amador, R., González, G., Ramírez-Umaña, C. J., Díez, B., & Pedrós-Alió, C. (2019). The influence of temperature and pH on bacterial community composition of microbial mats in hot springs from Costa Rica. MicrobiologyOpen, 8(10), e893. 10.1002/mbo3.893

Vos, M., Wolf, A. B., Jennings, S. J., & Kowalchuk, G. A. (2013). Micro-scale determinants of bacterial diversity in soil. FEMS Microbiology Reviews, 37(6), 936–954. 10.1111/1574-6976.12023

Walther, K. (2013). Ecology of Alkaline Hot Springs: Measuring Diversity and Structure of Chemosynthetic Communities. Dissertation, Notre Dame University.

Widder, S., Allen, R. J., Pfeiffer, T., Curtis, T. P., Wiuf, C., Sloan, W. T., Cordero, O. X., Brown, S. P., Momeni, B., Shou, W., Kettle, H., Flint, H. J., Haas, A. F., Laroche, B., Kreft, J.-U., Rainey, P. B., Freilich, S., Schuster, S., Milferstedt, K., … Soyer, O. S. (2016). Challenges in microbial ecology: Building predictive understanding of community function and dynamics. The ISME Journal, 10(11), 2557–2568. 10.1038/ismej.2016.45

Wiens, J. A. (1989). Spatial Scaling in Ecology. Functional Ecology, 3(4), 385. 10.2307/2389612

Xue, R., Zhao, K., Yu, X., Stirling, E., Liu, S., Ye, S., Ma, B., & Xu, J. (2021). Deciphering sample size effect on microbial biogeographic patterns and community assembly processes at centimeter scale. Soil Biology and Biochemistry, 156, 108218. 10.1016/j.soilbio.2021.108218

Yan, H., Yang, F., Gao, J., Peng, Z., & Chen, W. (2019). Subsoil microbial community responses to air exposure and legume growth depend on soil properties across different depths. Scientific Reports, 9(1), 18536. 10.1038/s41598-019-55089-8

Youssef, N. H., Couger, M. B., & Elshahed, M. S. (2010). Fine-Scale Bacterial Beta Diversity within a Complex Ecosystem (Zodletone Spring, OK, USA): The Role of the Rare Biosphere. PLoS ONE, 5(8), e12414. 10.1371/journal.pone.0012414

Zhou, J. (2009). Predictive microbial ecology. Microbial Biotechnology, 2(2), 154–156. 10.1111/j.1751-7915.2009.00090_21.x

