## Supplementary Material for "Fine-scale spatial patterns in hot springs mat bacterial communities"

**Supplementary Methods**

**DNA extraction, library preparation, and sequencing.**

Extraction of nucleic acids from samples was done as described by (Mackenzie et al., 2013). Briefly, samples were thawed and approximately 200 µl of the microbial mat was bead-beaten for rapid and thorough homogenization. Nucleic acids were extracted using a modified phenol: chloroform: IAA protocol and purified using Amicon Ultra-15 (Millipore, MA, USA). 16S Ribosomal DNA libraries were prepared following the procedure described by (Preheim et al., 2013). Succinctly, the 16S rRNA gene was amplified using as few cycles as possible (Polz & Cavanaugh, 1998). This was achieved by carrying out QPCR to identify the amplification cycle threshold (Ct) of each sample and modifying the thermocycler program to the minimum number of cycles possible. This QPCR was carried out in a final volume of 25 ul containing 5 µl of 5x HF buffer, 250 µM of dNTPs, 0.3 µM of the primers PE_16S_V4_U515_F (5’ - ACA CGA CGC TCT TCC GAT CTY RYR GTG CCA GCM GCC GCG GTA A – 3’) and PE_16S_V4_E786_R (5’ – CGG CAT TCC TGC TGA ACC GCT CTT CCG ATC TGG ACT ACH VGG GTW TCT AAT), 0.5x SYBR Green I nucleic acid stain (InvitrogenTM), 2.5 U of Phusion® High-Fidelity DNA Polymerase (New England BioLabs Inc.), and 10 ng of template DNA. The amplification program consisted of an initial denaturing step at 98 ºC for 3 min followed by an amplification step of 45 cycles of 30 s at 98 ºC, 30 s at 52 ºC, and 30 s at 72 ºC, and a final extension of 5 min at 72 ºC. All samples were normalized to the least concentrated sample according to the QPCR results. Samples were diluted according to Df= 2n, where Df is the dilution factor and n is the difference in Ct between the Ct of the sample to dilute and the lowest Ct (15 cycles in this experiment). Normalized templates were cycled for 15 cycles (based on QPCR results) with the same conditions as the QPCR reaction without SYBR Green with 5 µl of normalized template DNA. Due to the low number of cycles used, this PCR was done in quadruplicate, and products were pooled after amplification. Pooled PCR products were purified by magnetic beads (AgenCourt® AMPure® XP, Beckman Coulter). Each 100 µl of PCR was mixed with 85.5 µl of beads and left for 13 min to let the DNA bind to the beads. Then, samples were incubated for 15 min on a magnet (SPRIplate® 96-Ring) and washed three times with 100 µl of ethanol 70%. After ethanol evaporation, the DNA was eluted by 40 µl of EB buffer (QIAGEN) by 7 min incubation for elution and 15 min on the magnet to separate DNA from the beads.

Specific barcodes were added to each sample during the 9th cycle of PCR, with primers that overlapped with the former primers amplified region. This PCR was carried out in a final volume of 25 ul containing 5 µl of 5x HF buffer, 250 µM of dNTPs, 0.3 µM of the primers PE_III_F (5’ – AAT GAT ACG GCG ACC ACC GAG ATC TAC ACT CTT TCC CTA CAC GAC GCT CTT CCG ATC T – 3’) and PE_III_001-096 (5’ – CAA GCA GAA GAC GGC ATA CGA GAT NNN NNN N CGG TCT CGG CAT TCC TGC TGA ACC GCT – 3’; NNNNNNN represents the specific barcode), 2.5 U of Phusion® High-Fidelity DNA Polymerase (New England BioLabs Inc.), and four microliters of the previous purified PCR as template. The amplification program consisted of an initial denaturing step at 98 ºC for 2 min followed by an amplification step of 9 cycles of 30 s at 98 ºC, 9 s at 70 ºC, and 30 s at 72 ºC, and a final extension of 2 min at 72 ºC. This PCR was also done in quadruplicates, pooled after amplification, and purified by AgenCourt® AMPure® XP magnetic beads as described above. Finally, the libraries were multiplexed for Illumina sequencing. The multiplexing ratios were estimated by QPCR with the Illumina sequencing primers. This QPCR was carried out in a final volume of 25 µl containing 12.5 µl of 2x QuantiTecmastermix, 0.2 µM of the primers PE_seq_F (5' – ACA CTC TTT CCC TAC ACG ACG CTC TTC CGA TCT – 3') and PE_seq_R (5' – CGG TCT CGG CAT TCC TGC TGA ACC GCT CTT CCG ATC T – 3'), and 5 µl of the libraries as template. The amplification program consisted of an initial denaturing step at 95 ºC for 15 min followed by an amplification step of 45 cycles of 10 s at 95 ºC, 20 s at 60 ºC, and 30 s at 72 ºC, and a final extension of 5 min at 72 ºC. The samples were multiplexed and submitted for MiSeq Illumina sequencing at the BioMicro Center (MIT, Cambridge, MA). A set of 16S iTag sequences was obtained of 100 bp reads for the grid samples.

**Pre-processing and analysis of reads.**

The analysis of Illumina (MiSeq Sequencing System) sequences was performed first with Qiime (Caporaso et al., 2011) using its function split_libraries_fastq.py to demultiplex and quality filter sequences with the following parameters: the length of the barcode --barcode_type 7; max number of consecutive low-quality base -r 0; Q was used for quality --phred_quality_threshold 17. Next, demultiplexed and filtered reads were trimmed using MOTHUR (Schloss et al., 2011) to remove primers using the trim.seqs function and all reads were truncated at 77 and 81 bp for the forward and reverse read, respectively. This corresponded to positions 533-609 (forward) and 705-785 (reverse) of the Escherichia coli 16S rRNA sequence (Baker et al., 2003). After independently performing the previous steps on both forward and reverse reads, paired surviving reads were concatenated. The distribution-based clustering (DBC, (Preheim et al., 2013)) method was used to create OTUs from the concatenated sequences. DBC uses genetic distance and ecological information (i.e., the distribution of sequences across sampled environments) to inform the clustering algorithm. The goal of DBC is to accommodate differences at the level of genetic differentiation across taxa, reducing the number of redundant OTUs from sequences within the same population or created by sequencing error. The DBC algorithm uses read distribution information, the relative abundances of sequences within all samples, and genetic distance to inform clustering. Concatenated sequences were aligned to a modified SILVA bacterial alignment, which was trimmed to the same positions of the E. coli 16S rRNA sequence and concatenated in the same manner as the sequence data. Jukes-Cantor corrected distances were created using FastTree with the -makematrix option (Price et al., 2010) for both the aligned and unaligned sequences and used as input to the DBC algorithm. After creating OTUs with DBC, OTU representatives were defined as the most abundant sequence in the OTU. Non-16S rRNA sequences were removed from the final OTU list if the representative sequence alignment did not begin and end at the same positions as the reference alignment. Chimeric OTU representatives were identified with UCHIME 4.0 (Edgar et al., 2011) and removed. OTUs were assigned taxonomic information based on the representative sequence with the RDP classifier (Wang et al., 2007) using a bootstrap cut-off of 50% as recommended for sequences shorter than 250 bp.

The total number of 16S rRNA reads obtained after quality filtering and cleaning was 151,771 in the 86 samples. The number of reads varied considerably among samples, ranging between 271 and 5,591. We discarded 6 samples with less than 500 reads and normalized by rarefaction to 505 reads per sample before further analysis. After this process, we ended up with a total of 80 samples and 549 OTUs (of which 193 could not be assigned taxonomically, or 35.2%). The number of total OTUs found for Cahuelmó, Geyser, and Porcelana were 308, 374, and 185, respectively. A relatively small proportion of 15% (80 OTUs) was shared among all hot springs. Geyser contained 149 unique OTUs (27.5% of the total), the most diverse of the three hot springs, followed by Cahuelmó (124 unique OTUs, 22.6%), and Porcelana (36 unique OTUs, 7%)

**Supplementary Figures**

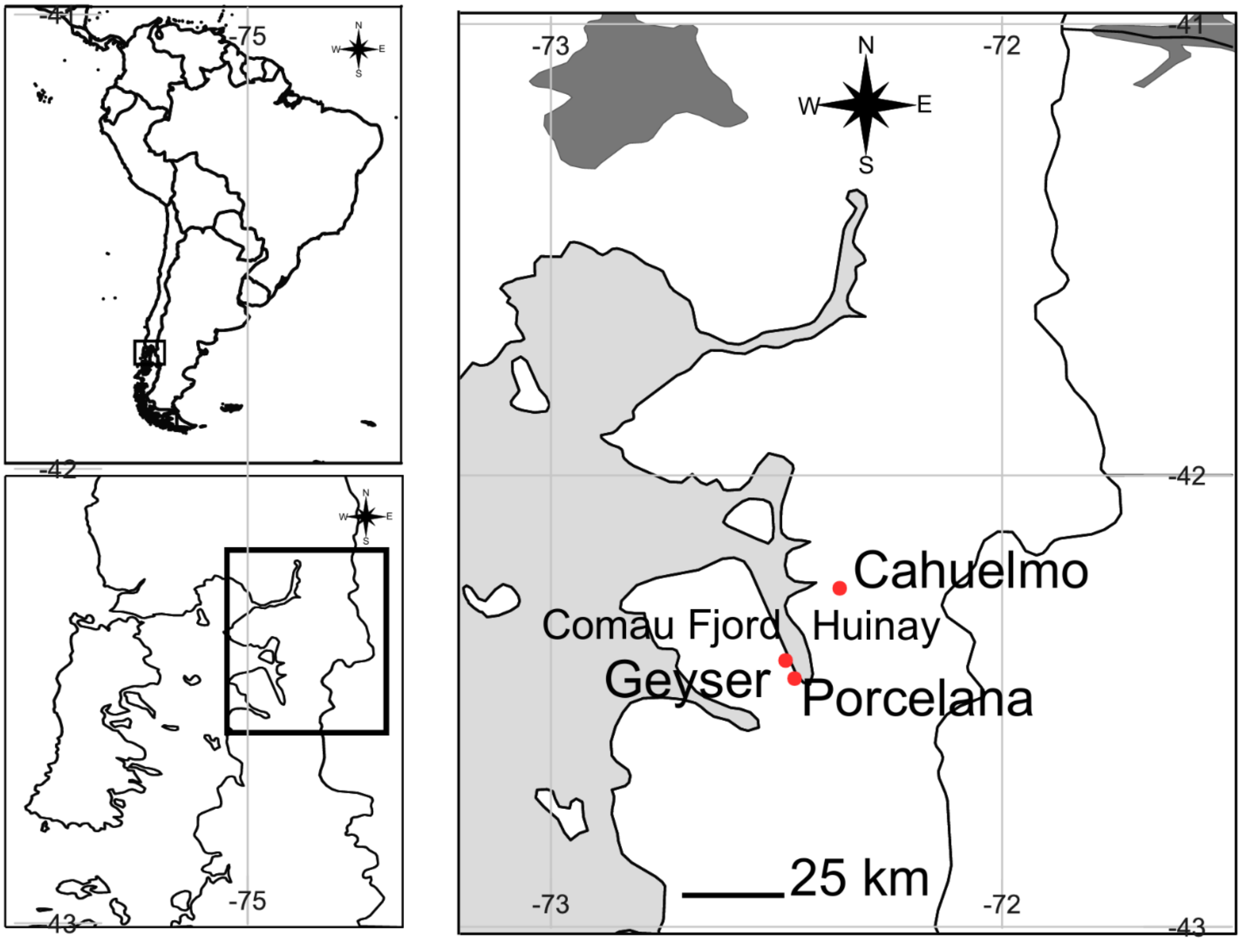

**Figure S1.** Map depicting the three localities studied.

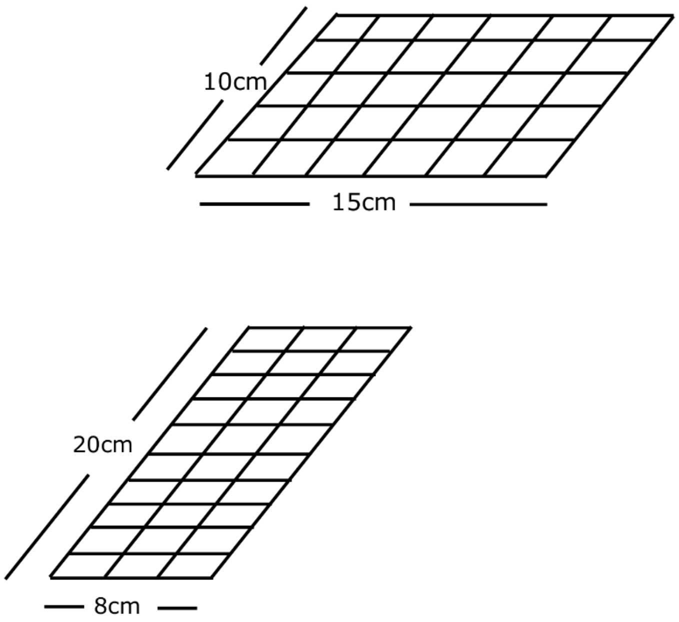

**Figure S2.** Sampling design for Cahuelmo (lower figure), Porcelana (upper figure), and Geyser sites (lower figure).

### References

Baker, G. C., Smith, J. J., & Cowan, D. A. (2003). Review and re-analysis of domain-specific 16S primers. *Journal of Microbiological Methods*, *55*(3), 541–555. https://doi.org/10.1016/j.mimet.2003.08.009

Caporaso, J. G., Lauber, C. L., Walters, W. A., Berg-Lyons, D., Lozupone, C. A., Turnbaugh, P. J., Fierer, N., & Knight, R. (2011). Global patterns of 16S rRNA diversity at a depth of millions of sequences per sample. *Proceedings of the National Academy of Sciences*, *108*(supplement_1), 4516–4522. https://doi.org/10.1073/pnas.1000080107

Edgar, R. C., Haas, B. J., Clemente, J. C., Quince, C., & Knight, R. (2011). UCHIME improves sensitivity and speed of chimera detection. *Bioinformatics*, *27*(16), 2194–2200. https://doi.org/10.1093/bioinformatics/btr381

Mackenzie, R., Pedrós-Alió, C., & Díez, B. (2013). Bacterial composition of microbial mats in hot springs in Northern Patagonia: Variations with seasons and temperature. *Extremophiles*, *17*(1), 123–136. https://doi.org/10.1007/s00792-012-0499-z

Polz, M. F., & Cavanaugh, C. M. (1998). Bias in Template-to-Product Ratios in Multitemplate PCR. *Applied and Environmental Microbiology*, *64*(10), 3724–3730. https://doi.org/10.1128/AEM.64.10.3724-3730.1998

Preheim, S. P., Perrotta, A. R., Martin-Platero, A. M., Gupta, A., & Alm, E. J. (2013). Distribution-Based Clustering: Using Ecology To Refine the Operational Taxonomic Unit. *Applied and Environmental Microbiology*, *79*(21), 6593–6603. https://doi.org/10.1128/AEM.00342-13

Price, M. N., Dehal, P. S., & Arkin, A. P. (2010). FastTree 2 – Approximately Maximum-Likelihood Trees for Large Alignments. *PLoS ONE*, *5*(3), e9490. https://doi.org/10.1371/journal.pone.0009490

Schloss, P. D., Gevers, D., & Westcott, S. L. (2011). Reducing the Effects of PCR Amplification and Sequencing Artifacts on 16S rRNA-Based Studies. *PLoS ONE*, *6*(12), e27310. https://doi.org/10.1371/journal.pone.0027310

Wang, Q., Garrity, G. M., Tiedje, J. M., & Cole, J. R. (2007). Naïve Bayesian Classifier for Rapid Assignment of rRNA Sequences into the New Bacterial Taxonomy. *Applied and Environmental Microbiology*, *73*(16), 5261–5267. https://doi.org/10.1128/AEM.00062-07
